# SENSE-PPI reconstructs protein-protein interactions of various complexities, within, across, and between species, with sequence-based evolutionary scale modeling and deep learning

**DOI:** 10.1101/2023.09.19.558413

**Authors:** Konstantin Volzhenin, Lucie Bittner, Alessandra Carbone

## Abstract

*Ab initio* computational reconstructions of protein-protein interaction (PPI) networks will provide invaluable insights on cellular systems, enabling the discovery of novel molecular interactions and elucidating biological mechanisms within and between organisms. Leveraging latest generation protein language models and recurrent neural networks, we present `SENSE-PPI`, a sequence-based deep learning model that efficiently reconstructs *ab initio* PPIs, distinguishing partners among tens of thousands of proteins and identifying specific interactions within functionally similar proteins. `SENSE-PPI` demonstrates high accuracy, limited training requirements, and versatility in cross-species predictions, even with non-model organisms and human-virus interactions. Its performance decreases for phylogenetically more distant model and non-model organisms, but signal alteration is very slow. `SENSE-PPI` is state-of-the-art, outperforming all existing methods. In this regard, it demonstrates the important role of parameters in protein language models. `SENSE-PPI` is very fast and can test 10,000 proteins against themselves in a matter of hours, enabling the reconstruction of genome-wide proteomes.

**Graphical abstract:** `SENSE-PPI` is a general deep learning architecture predicting protein-protein interactions of different complexities, between stable proteins, between stable and intrinsically disordered proteins, within a species, and between species. Trained on one species, it accurately predicts interactions and reconstructs complete specialized subnetworks for model and non-model organisms, and trained on human-virus interactions, it predicts human-virus interactions for new viruses.

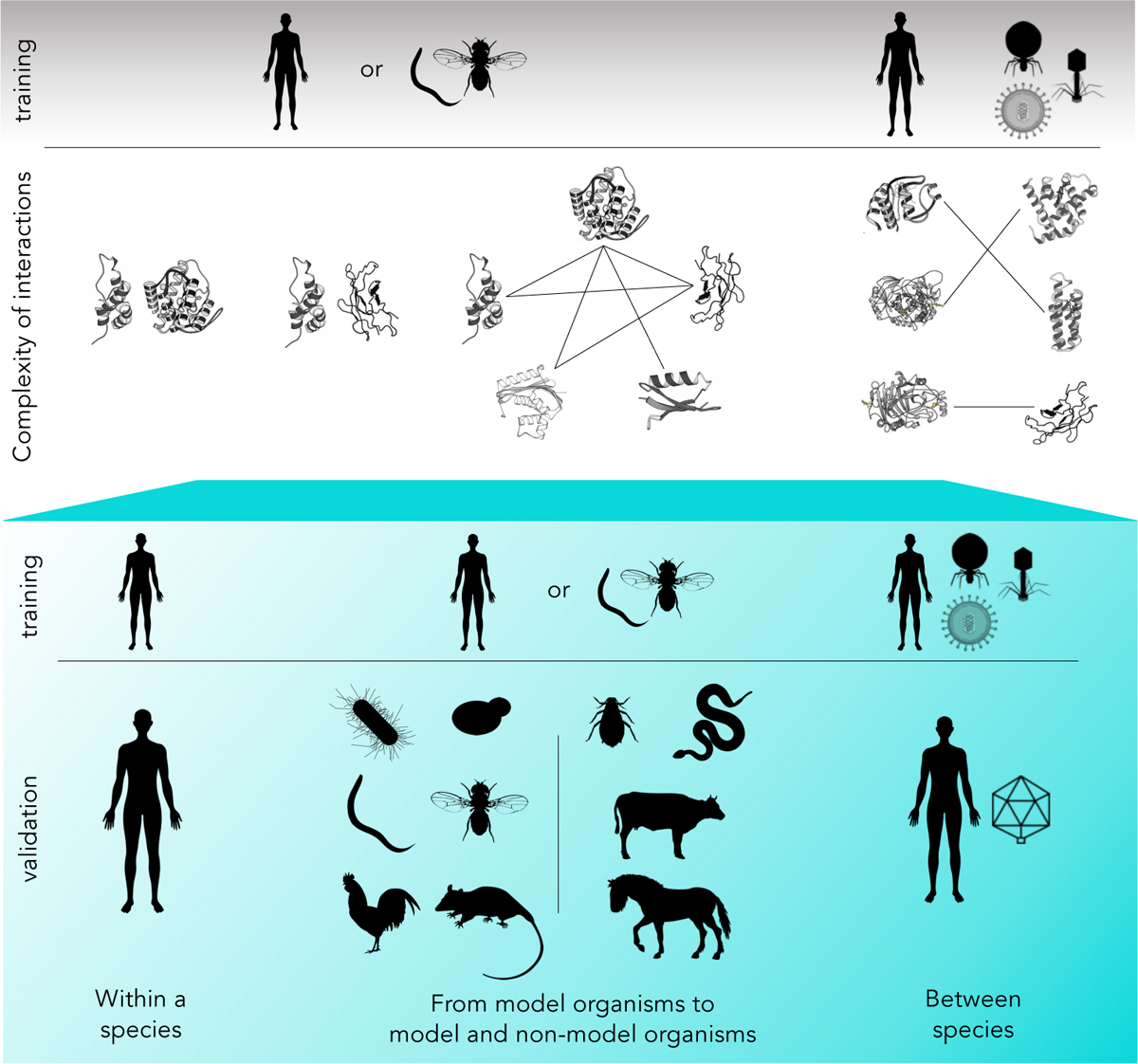

## 1 Introduction

Protein-protein interaction (PPI) networks play a key role in biology and medicine in the interpretation of protein functions in cellular processes. In the past two decades, working with networks has significantly advanced our understanding of the relationships between molecules [31, 59, 43, 34, 19, 20]. This was possible thanks to many computational attempts [37, 16, 29, 67, 49, 44, 12, 30, 64, 10, 27, 9, 13, 21], among which recent deep learning architectures [4, 61, 53, 52], and high throughput experimental methods such as yeast two-hybrid [62, 17] or tandem purification [2, 66] that have been extensively developed. A particular concern is their level of noise and incompleteness [22]. In addition, various studies give insights into the precise spatial organization and dynamic temporal remodeling of local protein interaction networks within the cell [48, 26] highlighting that PPI networks are only “projections” of particular spatio-temporal PPI realizations. For instance, within each individual, genomic alterations contribute to different PPI realizations. Accordingly, understanding any biological process demands defining three parameters: the composition of the “underlying protein network”, its organization in space, and its evolution over time. Future technological developments will bring an overwhelming amount of precise information on these genomics-spatio-temporal dimensions and sophisticated computational tools for extracting information from them are mandatory. Ultimately, PPI networks should be understood within biological frameworks including transcriptomic and epigenomic data. Here, we address the first step of this development, which is the construction of the “underlying protein network” which will serve as the basis for more sophisticated and realistic reconstructions. Because of the difficulties explained above, which are intrinsic to experimental data, *ab initio* computational reconstruction of PPIs is supposed to provide invaluable information leading to the identification of protein partners.

Today, the problem of PPI network reconstruction is becoming particularly important due to the wealth of sequence data available from different species and the need to understand proteomes within and between species which most of the time are non-model organisms, that is species that cannot grow in the laboratory, have a long life cycle, low fecundity or poor genetics for instance. At the same time, the identification of protein partners in PPIs puts new deep learning approaches to the test, for discovering sophisticated correlations within pairs of interacting protein sequences, and for estimating the absence of such correlations when the interaction does not exist in the cell. We take up this challenge and propose `SENSE-PPI`, an *ab initio* deep learning approach for protein partner identification, coupling layers of gated recurrent units (GRU) [5] with the ESM2 protein language model (PLM) encoding sequences [25], to achieve optimal identification for protein partners spanning a large spectrum of biological functions and leading to the reconstruction of PPI networks. Indeed, the complexity of the problem may vary depending on the features and the origins of proteins. `SENSE-PPI` is a generalizable architecture that can be applied to many types of protein datasets for PPI reconstruction.

It has been trained and tested on 1. the reference yeast dataset where the same proteins appear both in training and testing; 2. the human `D-SCRIPT` dataset [53] where close homologs might be shared in training and testing; 3. a second human dataset from the STRING database where sequences appearing in training and testing have been filtered based on homology; 4. the fly, warm, yeast, mouse, chicken and bacteria datasets used for validation based on training realized on a different species, in our case the human proteome; 5. the horse, cow, snake, and aphid datasets used for testing non-model organisms; 6. the human-virus dataset for protein interaction between species; 7. the dataset IDPpi [39] for interactions with human intrinsically disordered proteins. Based on this ample spectrum of datasets, we could evaluate the generalizability of `SENSE-PPI` and its limitations.

## 2 Results

We leverage the power of deep learning to identify correlations in interacting sequences, and to estimate their absence when there is no interaction. Our architecture, `SENSE-PPI`, exploits the latest generation of PLMs, ESM2, and recurrent NN, GRU, to predict the interaction of protein pairs with different features on a large scale. Validation is carried out using a wide range of interaction datasets of different complexities, from model and non-model organisms, inter-species interactions, human-virus interactions, and interactions with intrinsically disordered human proteins. A large new dataset of interactions in *Homo sapiens*, comprising over a 1 million interactions, is designed on the basis of a restrictive “neighboring-exclusion” condition that guarantees more accurate predictions, a property that is particularly required when laboratory tests are planned. SENSE-PPI is compared with state-of-the-art deep learning approaches, `PIPR`, `D-SCRIPT`, `Topsy-Turvy`, and `STEP`, in various scenarios.

### 2.1 A yeast dataset for a reference comparison

`SENSE-PPI` was first challenged on the Guo’s yeast dataset [12], used as a reference for method comparison. We compare `SENSE-PPI` with Guo’s performance, `PIPR` and `STEP`. All approaches provide significantly high scores, and these scores are very close to each other. The average Matthews’ correlation coefficient (MCC; see Methods) of `SENSE-PPI` is 95.46, which represents an improvement on the other deep learning models (94.77 and 94.17 for `STEP` and `PIPR` respectively), although this difference is not substantial (see Table SI 1). The exceptional success of all methods is explained by the characteristics of the computational experiment. In fact, all models were tested by performing 5-fold cross-validation on a dataset of protein pairs where the same protein appears in several pairs in the full dataset. Since the testing data represent a randomly chosen subset of sequence pairs at each cross-validating fold, in the validation step, each model mostly evaluates proteins already “seen” during training. Such pairs are known to show better performance than those excluded from training [35, 13].

### 2.2 Training on the human proteome and testing on model and non-model organisms

Recent global efforts to sequence the biodiversity of species [60, 3, 23, 24, 41], make PPI predictions in yet unexplored organisms a major challenge. Here, we test `SENSE-PPI` on non-model species and assess its generalization capabilities for predicting interactions in model organisms that are phylogenetically distant, to varying degrees, from the species used for training. The `SENSE-PPI` model was trained on *H. sapiens* and tested on PPI data from several different species: *H. sapiens, Mus musculus, Drosophila melanogaster, Caenorhabditis elegans, Saccharomices cerevisiae, Escherichia coli, Gallus gallus, Equus caballus, Bos taurus, Notechis scutatus*, and *Acyrthosiphon pisum*. The first seven organisms are model organisms but the last four are not.

#### 2.2.1 Benchmark testing for model organisms

To compare the performance of `SENSE-PPI` with state-of-the-art approaches on model organisms, we repeated the training and testing procedures described in [53]. `SENSE-PPI` was trained on the large `D-SCRIPT` human dataset, specially designed to avoid protein redundancies in training and validation sets. Table 1 presents the comparison of `SENSE-PPI` performance with those of `PIPR`, `D-SCRIPT` and `Topsy-Turvy` trained on the same human data. The overall performance remains high for all four DL models when testing is carried out within *H. sapiens*. However, scores decline progressively when tests are carried out on the other four model species. As the species’ evolutionary distance from *H. sapiens* increases, performance decreases progressively: the most accurate predictions are obtained for *M. musculus* and the worst for *S. cerevisiae*. `SENSE-PPI` outperforms all other DL methods on all model species tested and, more importantly, offers greater generalization ability in several respects. First, the ability of `SENSE-PPI` to distinguish between positive and negative interactions is measured with the AUROC score (see Methods) which remains above 0.9 for all test sets, while `D-SCRIPT` goes from an AUROC score of 0.833 for *H. sapiens* down to 0.790 for *S. cerevisiae*. Second, `SENSE-PPI`’s performance results in an F1-score (see Methods) of 0.848 within *H. sapiens* that remains above 0.737 for *M. musculus, D. melanogaster*, and *C. elegans*. This result shows that PPI networks of model species whose common ancestor is estimated at 708 million years from *H. sapiens* [18] remain well predicted by `SENSE-PPI` trained on *H. sapiens*. The `SENSE-PPI` F1-score decreases to 0.555 for *S. cerevisiae*, sharing a common ancestor with *H. sapiens* at 1,300 million years. It should be noted, however, that the performance of `SENSE-PPI` on this very distant species is superior in terms of F1-score to that of `PIPR`, with 0.555 versus 0.456 on *M. musculus*, and `D-SCRIPT`, with 0.402 on *H. sapiens*. Overall, `SENSE-PPI` significantly outperforms previously developed methods on both single-species and cross-species tests.

**Table 1:** Evaluation of SENSE-PPI trained on the human D-SCRIPT dataset. Training of `SENSE-PPI` is carried out on the human `D-SCRIPT` dataset. Testing was carried out on species not present in the training data: mouse, fly, worm, and yeast (top). Testing on the human dataset is also shown (bottom). The best values for every specific metric and dataset are highlighted in bold. Scores for `PIPR`, `D-SCRIPT` and `Topsy-Turvy` in the first 4 test sets were taken from [52]. The scores for `PIPR` and `D-SCRIPT` in the human test set were taken from [53]. The evaluation of `Topsy-Turvy` on human data was recomputed.

| Species | Model | AUPRC | AUROC | F1-Score |
| --- | --- | --- | --- | --- |
| *M. musculus* | PIPR | 0.526 | 0.839 | 0.456 |
|  | D-SCRIPT | 0.663 | 0.901 | – |
|  | Topsy-Turvy | 0.735 | 0.934 | – |
|  | SENSE-PPI | **0.859** | **0.973** | **0.782** |
| *D. melanogaster* | PIPR | 0.278 | 0.728 | 0.196 |
|  | D-SCRIPT | 0.605 | 0.890 | – |
|  | Topsy-Turvy | 0.713 | 0.921 | – |
|  | SENSE-PPI | **0.847** | **0.969** | **0.742** |
| *C. elegans* | PIPR | 0.346 | 0.757 | 0.235 |
|  | D-SCRIPT | 0.550 | 0.853 | – |
|  | Topsy-Turvy | 0.700 | 0.906 | – |
|  | SENSE-PPI | **0.821** | **0.963** | **0.737** |
| *S. cerevisiae* | PIPR | 0.230 | 0.718 | 0.140 |
|  | D-SCRIPT | 0.399 | 0.790 | – |
|  | Topsy-Turvy | 0.534 | 0.850 | – |
|  | SENSE-PPI | **0.657** | **0.914** | **0.555** |
| *H. sapiens* | PIPR | 0.835 | 0.960 | 0.763 |
|  | D-SCRIPT | 0.516 | 0.833 | 0.402 |
|  | Topsy-Turvy | 0.703 | 0.895 | 0.711 |
|  | SENSE-PPI | **0.917** | **0.984** | **0.848** |

#### 2.2.2 New human training dataset improves accuracy

Motivated by the design of a fair training dataset on which to evaluate `SENSE-PPI`, we constructed the `SENSE-PPI` human dataset from interactions in the STRING database v11.5. The set of negative protein pairs was defined by satisfying a stringent “neighboring-exclusion” condition ensuring that A-B is a negative interaction in the training set if: 1. A and B are not known to interact in STRING, 2. no homolog of A is known to interact with a homolog of B at more than 40% sequence identity in STRING, and 3. no homolog, at more than 40% sequence identity, of a known interactor of B is known to interact with a homolog of A. Note that the first two conditions are also used in the construction of the `D-SCRIPT` human dataset. This dataset is double the size of the `D-SCRIPT` human dataset, and contains more than 86,000 positive and 860,000 negative pairs. We have trained `SENSE-PPI` on this new dataset and have compared `SENSE-PPI` performance on the four `D-SCRIPT` test sets from four different species [53]. Table 2 shows the F1-score, precision, and recall. Even though the `SENSE-PPI` human dataset leads to better performance (F1-score) in 3 cases out of 4, the increase remains on par. For practical usage, it is important to note that the trained models provide different precision/recall ratios. In particular, the `SENSE-PPI` human dataset obtains higher values of precision (trading off some recall instead) and might be more valuable when lab experiments should be conducted and target partners identified. It will produce fewer false positives thus increasing the proportion of relevant interactions among all positive predictions.

**Table 2:**
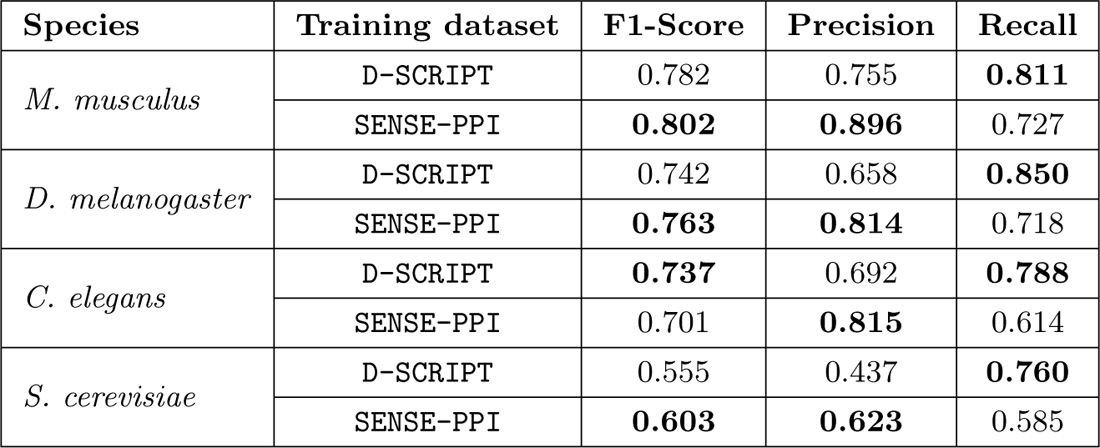
Evaluation of SENSE-PPI trained on two human interaction datasets. Training was carried out on the `SENSE-PPI` and `D-SCRIPT` datasets. Testing was carried out on the four model species in [53]. For each species and each metric, the best performances are highlighted in bold.

#### 2.2.3 Testing across the evolutionary tree on model and non-model species

Furthermore, we asked whether `SENSE-PPI` behavior is preserved across the phylogenetic tree and, in particular, for non-model species. For this, we constructed `SENSE-PPI` datasets for the four non-model and six model species (see Methods). A coherent behavior of `SENSE-PPI` across species and time evolution would support the use of `SENSE-PPI` on species at a fixed evolutionary distance from the training one for which little is known about protein interactions. Consistent with our expectations, the behavior of `SENSE-PPI` is illustrated in Figure 1 (see also Table SI 4).

**Figure 1:**
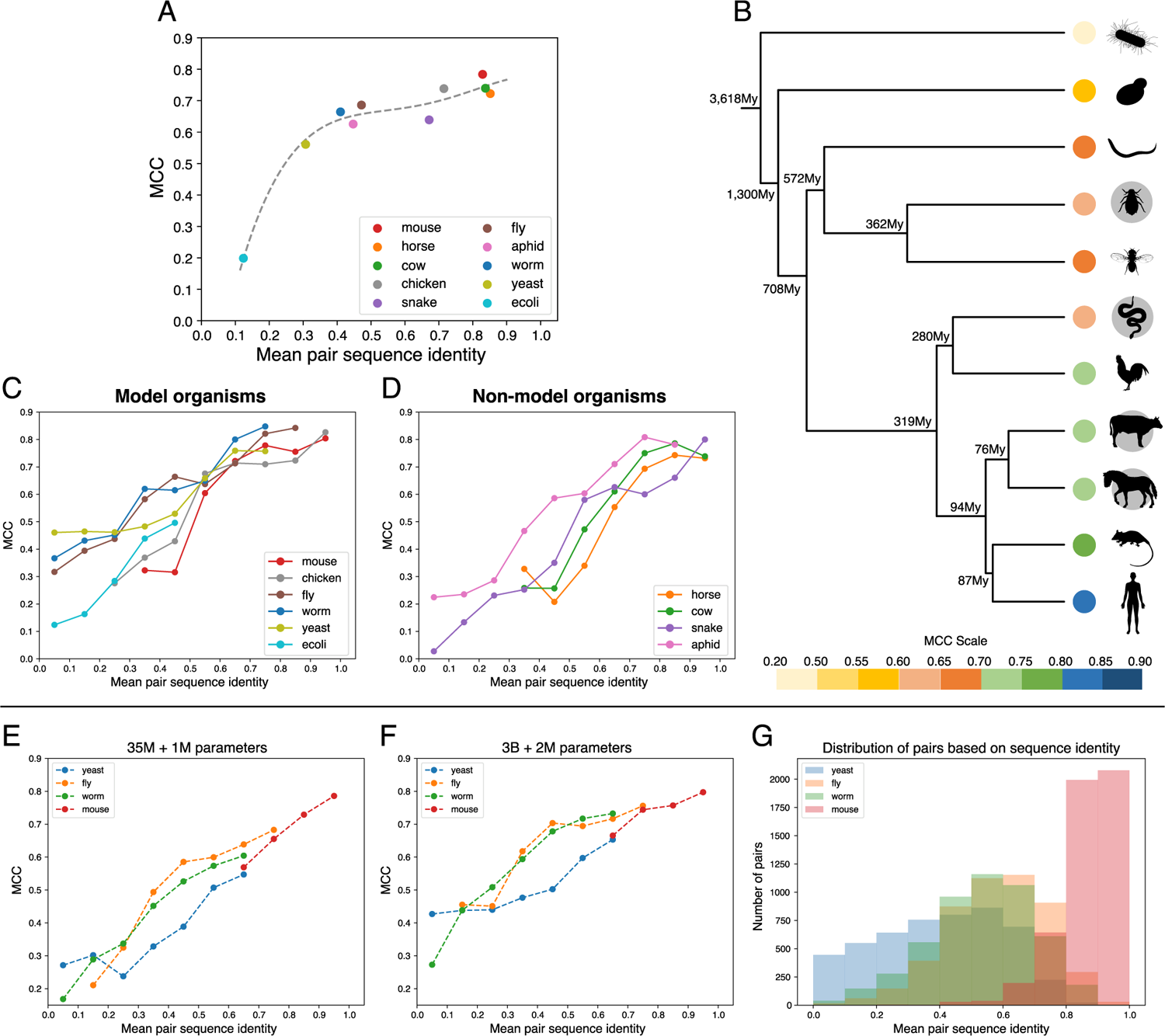
Performance of SENSE-PPI on six model and four non-model organisms. `SENSE-PPI` is trained on the `SENSE-PPI` human dataset. For each experiment, the testing dataset comprises 5,000 positive and 50,000 negative interactions. **A**: Matthews correlation coefficient (MCC) versus mean pair sequence identity (see Methods) for the ten experiments. Dotted line: least squares polynomial fit with degree 4. **B**: Phylogenetic tree of species used for `SENSE-PPI` testing and performance evaluation. The color scale, labeling the species in the tree, corresponds to the MCC value obtained in testing (as reported in A); colors go from blue (high MCC) to yellow (low). Non-model species are shown with a grey disk behind them. The tree was reconstructed using phyloT v2 (http://phylot.biobyte.de/). Evolutionary information on species divergence times was obtained with TimeTree [18]. F1 and MCC scores are reported in Table SI 4. The MCC score for *H. sapiens* trained on *H. sapiens* is 0.836. **C and D**: Plots of MCC scores versus mean pair sequence identity for model (C) and non-model (D) organisms. **E and F**: Comparison of two versions of `SENSE-PPI` differing by the size of the ESM2 embedding module. The trend of `SENSE-PPI` MCC scores versus the mean pair sequence identity is shown for 4 different testing datasets coming from [53] for the “light” version (E) and the regular version (F). **G**: Distribution of pairs in four test sets for panels E-F based on the mean pair sequence identity. The data on panels C-G are presented in bins of 10% across the values of mean pair sequence identity. The MCC on panels C-F is computed for a given bin only if it contains at least 40 positive and 400 negative samples.

The evaluation of `SENSE-PPI` on 10 testing datasets across species provides valuable insights, although it does not offer a comprehensive assessment of performance. Hence, to tackle the redundancy problem inherent in the input data seen in training [35, 13], we systematically filtered the proteins in the test sets according to different levels of sequence identity from proteins in the training set and evaluated `SENSE-PPI` precision accordingly. Previous attempts to analyze the effects of the proximity of training and testing sets mostly used the concepts of C1/C2/C3 classes [35, 13]. These classes divide the testing set into three subgroups based on how many proteins in a pair were already present in training (two, one, or none respectively). In this work, however, we focus on how performance varies with the increase in the evolutionary distance between the tested species and the species from the training data without relying on this discrete division into three classes.

For this, different extents of sequence identity were measured with MMseqs2 [54] by searching for the closest matches (i.e., consecutive k-mer matches, based on sequence identity without gaps) between testing and training sequences. For each protein pair in the testing set, we computed a so-called “mean pair sequence identity”: first, we computed the maximum sequence identities for both proteins in a pair with respect to all proteins in the training set, and then, we considered the mean of these two values (see Methods). Figure 1A shows the distribution of MCC scores with respect to the mean pair sequence identity for ten different species. We observe a clear positive correlation: the larger the mean pair sequence identity is, the better the performance. This agrees well with the evolutionary distance between species (Figure 1B): species that are closer to the training data (*H. sapiens*) tend to have higher scores. However, one can observe a tendency of non-model organisms to have slightly lower scores: Figures 1CD show the MCC vs. pair sequence identity plots for model and non-model organisms respectively. Here, instead of simply taking the mean pair sequence identity for all test entries, the data were divided into bins of size 0.1. The two plots show that the main difference in the evaluation of model versus non-model organisms concerns low values of mean pair sequence identity. With the only exception of *E. coli* (which is the farthest organism from the training data in terms of evolutionary distance), the MCC scores for model organisms are not lower than 0.3. At the same time, non-model organisms start to have scores greater than 0.3 only when the mean pair sequence identity rises beyond 40%.

As a proof of concept, we tested the non-model organism *A. pisum* after training `SENSE-PPI` on a dataset made of species that are evolutionary closer to aphids than to humans. This dataset is a merge of two separate datasets for *D. melanogaster* and *C. elegans* (see Methods), sharing a common ancestor with the aphid at 362My and 572My respectively compared to the common ancestor shared with *H. sapiens* at 708Mys. This merge was necessary due to the absence of sufficiently many high-confidence interactions in *D. melanogaster* alone. Hence, a relatively close model species was chosen, *C. elegans*, so that the size of the resulting testing set could be comparable to the human `D-SCRIPT` dataset used in Table 1. In line with our expectations, PPI predictions on non-model organisms are improved when closer model organisms are used to train `SENSE-PPI`. Test scores for *A. pisum* improved from 0.660 to 0.701 in F1-score, and from 0.626 to 0.689 in MCC. Moreover, by combining the training set from the two species with the `SENSE-PPI` human dataset, we can further improve predictions, achieving an F1-score of 0.742 and an MCC of 0.724. This observation should be taken as a general recommendation, even though the improvement is not exceptionally high due to `SENSE-PPI` general high performance over a large interval of mean pair sequence identity values as illustrated by the plateau of the curve in Figure 1A.

#### 2.2.4 Larger protein language models improve performance

PLMs significantly contribute to enhanced performance. To see this, we consider two sizes of ESM2 embedding modules and analyze how MCC scores change at different values of the mean pair sequence identity for the four `D-SCRIPT` datasets of *M. musculus, D. melanogaster, C. elegans*, and *S. cerevisiae* (Figure 1EF). Figure 1E shows the `SENSE-PPI` performance obtained with the smaller version of ESM2, based on 35M parameters (with 1M more parameters used in the rest of the `SENSE-PPI` architecture), and Figure 1F demonstrates the performance of the `SENSE-PPI` regular version, based on 3B parameters (with 2M more parameters for the rest of the `SENSE-PPI` architecture). The ESM2 model with 3B parameters highly improves the predictions on pairs of proteins with low values of mean pair sequence identity compared to the smaller version of ESM2. For instance, for a pair sequence identity *<* 0.2, predictions on the fly display an MCC of 0.2 (Figure 1E) versus 0.4 (Figure 1F). Moreover, for entries with high values of mean pair sequence identity, the performance based on both embeddings remains comparable. This remains true for all 4 datasets and shows that ESM2 based on a higher number of parameters has better generalization capability.

### 2.3 Evaluation on the human intrinsically disordered protein dataset

Approximately 33% of eukaryotic proteins have significant disordered regions, with an increasing occurrence of disorder in higher organisms [38], and predicting interactions between proteins that are possibly disordered becomes an urgent demand [32, 63, 50, 51]. Deep learning approaches based on sequences are expected to open the way to this challenge. `SENSE-PPI` was additionally compared to the IDPpi model [39] designed to predict the interaction between proteins with stable structure and intrinsically disordered proteins (IDPs). Table SI 2 shows the results for both models. The models were tested on the same five test sets used in [39] but were trained differently. Whereas the IDPpi training set included the IDPs in the test data, `SENSE-PPI` was trained on the `D-SCRIPT` human dataset, which contained no information on IDPs. The same performances obtained by the two approaches suggest that by providing more extensive training (note that the `D-SCRIPT` dataset is 20 times larger than the IDPip training dataset), `SENSE-PPI` can obtain relatively accurate predictions on IDP interactions. This test once again highlights the generalization capabilities of `SENSE-PPI` enabling us to extend its use to process IDPs with fairly good accuracy.

### 2.4 Human-virus PPIs

Predicting and understanding virus-host PPIs is important for developing new therapeutic interventions, but knowledge of virus-host interactions covered by current databases, such as VirHostNet [11], is limited. We therefore tested the ability of `SENSE-PPI` to predict interactions between a known human protein and an unknown viral protein. We trained `SENSE-PPI` on a human-virus PPI dataset [61] that contains interactions between human and viral proteins, i.e. only cross-species interactions are included. Data were split between training and testing, so that test data only included interactions for Epstein-Barr and influenza viruses. The model was trained on the set with a large variety of viral species interacting with human proteins (for dataset details see Methods). We required that all human proteins involved in the test have been seen during training (this dataset corresponds to class “C2” in [13]), a condition that is reasonable in terms of potential applications. The results of this test are presented in Table SI 3. The overall high scores may be explained by the fact that the model is already familiar with all the human proteins present in the data [13]. However, it should be noted that the viral proteins from training and testing are drastically different: by searching the closest matches for every test viral protein in the training set (using MMseqs2, see Methods), we computed a mean sequence identity of 0.21 for the Epstein-Barr virus and of 0 for the Influenza viruses (no matches were found even with sensitivity parameter values in the range of 7-10 [54]). Nonetheless, the protocol can be potentially useful for further exploration of interactions between different species, where the model is familiar with all the target proteins of one of the organisms of interest. The problem of predicting cross-species interactions between two unknown proteins remains wide open.

### 2.5 PPI network reconstruction

In order to verify the performance of `SENSE-PPI` on the *ab initio* reconstruction of PPI networks, we carried out several tests to reconstruct PPI networks for sets of a few dozen proteins from the STRING database (version 11.5) [58]. For each test, we set one or two proteins as “seeds” and collected known partners on the basis of high confidence for physical interaction according to STRING. We then calculated `SENSE-PPI` predictions for a comparative analysis with STRING data. Figures 2 and 3 show four examples of these tests chosen to illustrate the general characteristics of `SENSE-PPI` behavior: 1. False positives are frequently related to indirect interactions, that is non-physical interactions with proteins sharing a common partner in the network (Figure 2A); 2. `SENSE-PPI` often misses interactions for weakly connected proteins (Figure 2B); 3. it successfully identifies interactions involved in functionally distinguished subnetworks (Figure 2A and Figure 3); 4. it successfully predicts PPIs in other species (Figure 2C). These four sets group interacting human proteins (Figure 2AB), *C. elegans* proteins (Figure 2C) and *M. musculus* proteins (Figure 3). The tests were carried out on the model trained on the `D-SCRIPT` human dataset.

**Figure 2:**
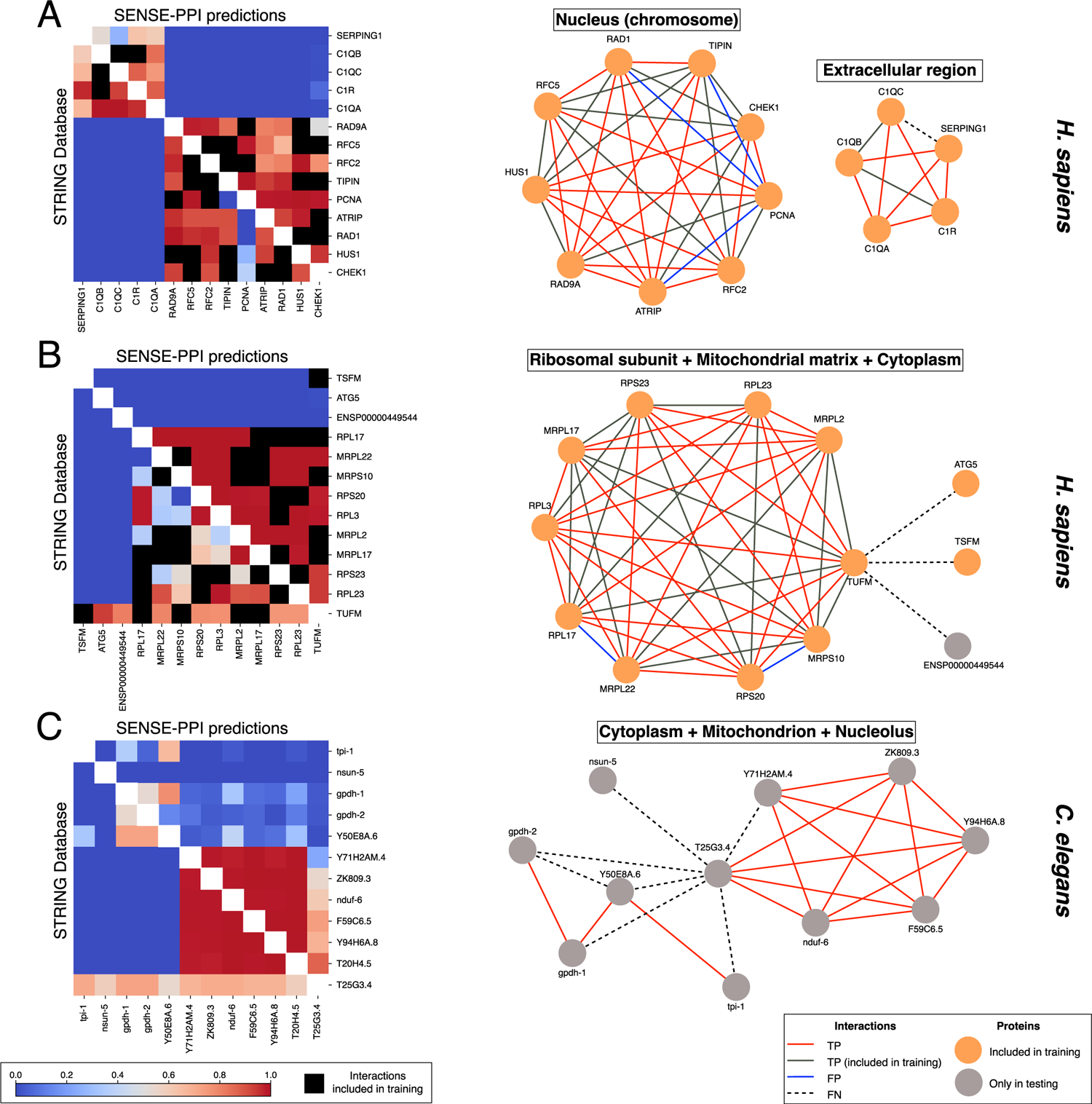
SENSE-PPI *ab initio* reconstruction of PPI networks and comparison with data from the STRING database. The three sets of potential protein partners are seeded on the C1R and RFC5 proteins in *H. sapiens* (A), the TUFM protein in *H. sapiens* (B), and the T25G3.4 protein in *C. elegans* (C). Each network is described by a heatmap (left) and a graph of interaction (right), where nodes are proteins and edges are interactions. The bottom triangle of the heatmap reports the STRING confidence scores (v11.5), while the upper triangle describes `SENSE-PPI` prediction scores. Positive and negative interactions present in training/validation are omitted from testing (black masked cells). The graphs represent predicted connections and trace (with distinct colors) information coming from both the STRING database and the training set. Nodes represent proteins present at least once in training (orange), and proteins present only in testing (grey). Four kinds of interactions are essential for comparison: interactions found in STRING and identified by `SENSE-PPI` (TP), interactions identified by `SENSE-PPI` that are not present in STRING (FP), interactions that are not found by `SENSE-PPI` but are present in STRING (FN), and interactions not found in STRING and not detected by `SENSE-PPI` (TN). Hence, edges show TP present in training (grey) and absent in training (red), FP (blue), and FN (dashed black). The absence of an edge indicates a TN. The `SENSE-PPI` prediction threshold used for graph reconstruction is 0.5, for the three networks.

**Figure 3:**
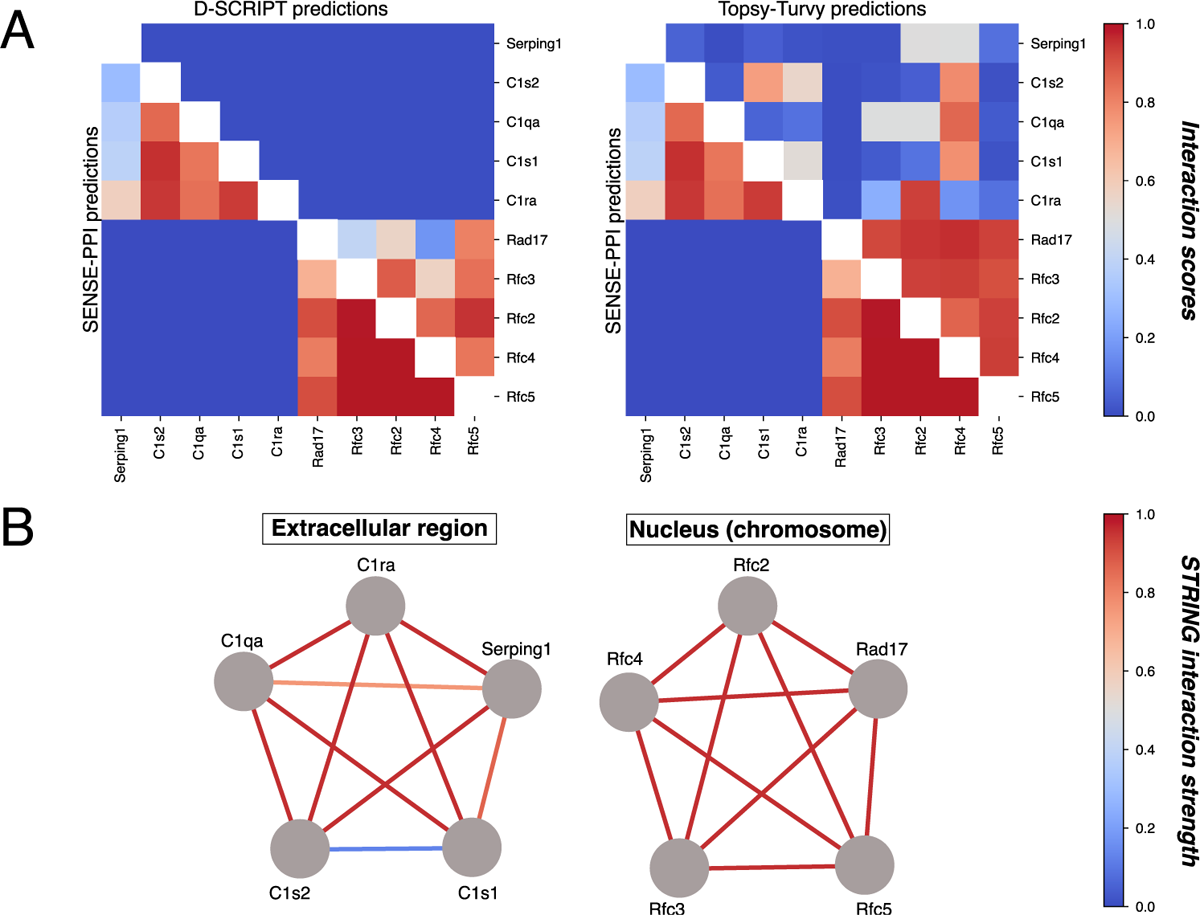
PPI network reconstruction seeded on the protein complexes C1-Serping1 and RFC-Rad17 in *M. musculus*. A. `SENSE-PPI` trained on the `D-SCRIPT` human dataset was used to reconstruct two networks with independent functionality in the cell. None of the proteins in the sets were observed during training, hence the full set of predicted interactions was considered in the evaluation. The heatmaps show `SENSE-PPI` (lower triangle), `D-SCRIPT` (upper triangle, left), and `Topsy-Turvy` (upper triangle, right) predicted interaction scores, for all possible protein pairs in the set. According to STRING v11.5, all RFC-Rad17 partners and C1-Serping1 partners physically interact with each other while no interaction between these two groups of proteins is known (see B). B. Two graphs representing STRING data on the two complexes. No mouse protein (gray) has been seen in training which is done on *H. sapiens*. Edge colors represent the STRING confidence score for the interactions. Scale colors as in Figure 2.

Three sets of proteins are analyzed in Figure 2 where heatmaps display confidence scores for STRING (lower triangle) and prediction scores for `SENSE-PPI` (upper triangle). Although we used the same color spectrum for both sources, there is no direct correlation between the `SENSE-PPI` scale and the STRING scale, since our model was trained on binary labels and STRING provides confidence estimates that were not taken into account during `SENSE-PPI` training (except for the fact that we only used STRING high-scoring interactions in the training set). In fact, interactions used for training or validation were not used for testing (see black cells in heatmaps). An intuitive representation of the heatmap data is illustrated by three associated graphs. The first protein set (Figure 2A) is seeded on C1R and RFC5 proteins from *H. sapiens*, and encompasses 12 other human proteins meant to interact with at least one seed protein. Proteins for which the majority of positive interactions in a given subset were already present during training have been omitted. C1R and RFC5 were chosen based on their subcellular localization: C1R is secreted in extracellular regions, while RFC5 is expressed in the nucleus. Moreover, these proteins possess different functional characteristics: while C1R is the first component of the classical pathway of the complement system within the immune system, RFC5 takes part in DNA replication in the cell cycle. `SENSE-PPI` clearly distinguishes the two functionally independent networks without any uncertainty, assigning very low scores to all protein pairs across the subnetworks. Some false positive interactions are present and strictly concentrated in the RFC5 complex. They concern the PCNA protein, which interacts, at different strengths in STRING, with RFC5 and many, but not all, of its partners. For this set, `SENSE-PPI` inference of false positives is often observed for STRING highly connected proteins. In such cases, reconstructions are not completely false, as a considerable proportion of errors are made on protein pairs that may not interact physically, but are either functionally linked or homologous to actual physical partners.

The second set is seeded on the human TUFM protein and 12 other proteins (Figure 2B), known to belong to the TUFM complex as described in STRING with the highest scores. TUFM is an elongation factor localized in the mitochondrial outer membrane. `SENSE-PPI` shows an excellent reconstruction ability for this complex, with very high scores provided for the interactions even though multiple proteins have not been seen in training. However, a weakness is observed for proteins with a low degree of interaction in the network, such as TSFM, ATG5, and ENSP00000449544, where `SENSE-PPI` fails to predict interaction. Indeed, `SENSE-PPI` tends to be conservative and does not assign positive labels to potential interactions, resulting in a considerable number of false negatives especially those associated with low-degree nodes.

The third set is seeded on the T25G3.4 protein from *C. elegans* (Figure 2C), a protein localized in the mitochondrial inner membrane. `SENSE-PPI` is able to address the problem of an *ab initio* network reconstruction in other species relatively well. As before, we observe that high-degree nodes give rise to fewer errors, while other parts of the network are more difficult to predict. The heatmap clearly shows that while `SENSE-PPI` reliably predicts a complete subnetwork, the remaining interactions were predicted with lower scores.

It is important to point out that even though we use STRING both for training and final evaluation, the testing examples may contain low-scoring interactions, indicating that some parts of this information cannot be taken as ground truth and should be treated with caution.

To further check the ability of `SENSE-PPI` to correctly infer interactions in other organisms, we reconsidered as seeds the C1 complex (C1r, C1s, C1q) with its plasma protease C1 inhibitor Serping1 playing a role in the immune response, and the replication factor RFC complex with Rad17, forming a DNA damage checkpoint complex in the cell cycle, both in *M. musculus*. In the mouse, as in human, these two subnetworks are not supposed to share common partners since the two complexes involved assure drastic differences in functions and subcellular localizations. We compared `SENSE-PPI` to the `D-SCRIPT`and `Topsy-Turvy` models [52] (see also Table 1), where training was performed for all models on the human `D-SCRIPT` dataset. For each seed protein, we chose four partners with the highest confidence score in STRING. Figure 3 shows two heatmaps where network reconstruction shows two separate clusters in `SENSE-PPI` predictions (lower triangle). `D-SCRIPT` (upper triangle, Figure 3A) correctly predicts the RFC-Rad17 complex network but fails on the C1-Serping1 complex. `Topsy-Turvy` improves `D-SCRIPT` predictions on the scoring values of interaction pairs in the RFC-Rad17 complex by being more confident in the predictions. On the C1-Serping1 complex, it identifies some interactions with variable confidence compared to `D-SCRIPT` but much less sharply than `SENSE-PPI`. With an increased number of true positives, the number of false positives increases as well though, and the model incorrectly finds nonexistent interactions between proteins belonging to different complexes. In contrast, with a clear separation between C1-Serping1 and RFC-Rad17 complexes, `SENSE-PPI` shows the quality of its predictions.

## 3 Discussion

Current PPI experimental datasets are still far away from having reached completion, even on well-studied model organisms such as yeast or humans [55, 14], and their curation is an active area of research [46]. Today, physical interaction networks obtained by high-throughput techniques are found to include numerous nonfunctional PPI [22] and at the same time, many missing true interactions. This is the main reason that an *ab initio* computational reconstruction of PPIs would provide invaluable information.

We introduced `SENSE-PPI`, a deep learning model for predicting protein-protein interactions based entirely on sequence information. Trained in a single species, `SENSE-PPI` predicts the proteome within the same species and generalizes to other species. Trained on physical interactions between proteins with stable structures, it generalizes to proteins interacting with IDPs. Trained on the interaction between proteins of human and viral species, it generalizes to the interaction between human proteins and proteins of new viruses.

The high performance of `SENSE-PPI` in single model species predictions has been demonstrated here to open up new possibilities in PPI network reconstruction for non-model organisms, which are often characterized by very sparse or absent biological information. Our analysis shows that training on a model species such as *H. sapiens* and testing on a phylogenetically distant species such as *C. elegans*, yet provides excellent inference. Moreover, we show that if we construct a training dataset out of model organisms that are evolutionary close to a non-model species, we are able to improve the performance: we succeeded in improving the test scores on *A. pisum* by switching the training data from *H. sapiens* to a combination of *C. elegans* and *D. melanogaster*. As a general PPI reconstruction strategy for a non-model organism, the user must first identify the best model organism(s) close to the non-model species, train on it, and infer PPI interactions in the non-model species.

`SENSE-PPI` is fast, exceeding the time limits of other DL approaches and outperforming the particularly time-consuming docking approaches [27]. The major advantage of ESM2, used in `SENSE-PPI`, over other PLM models is that it is much faster (several minutes versus several hours compared to ProtT5-XL-UniRef50 [8] for example) while providing highly informative matrices. Indeed, ESMFold, which works with a similar embedding, was able to compete with AlphaFold in terms of performance, being 60 times faster and enabling the reconstruction of over 600 million protein structural models [25]. The advantage of deep learning architectures over other computational attempts in PPI reconstruction is that training concentrates all the computational weight, while the combinatorics of the problem (i.e. the quadratic number of potential pairwise interactions among the proteins) is handled efficiently by the trained architecture. As expected, `SENSE-PPI` shows that the computational issue can be overcome, and that PPIs of tens of thousands of proteins can now be handled. Indeed, the identification of protein partners in a 100-protein dataset takes to takes `SENSE-PPI` around 2 minutes with a single NVIDIA GeForce RTX 3090. Given that the computational time for ESM2 embeddings scales linearly in the number of proteins (the tensors are computed only once for each protein and then reused) and the prediction time scales quadratically, we can can estimate that it takes around 2 hours to predict the partners of a set of 1,000 proteins and around 3 to 4 hours for 10,000.

Another major advantage of `SENSE-PPI` over other structure-based approaches to protein interactions, including docking, is that `SENSE-PPI` achieves far better results based solely on sequences, opening up new directions for development. The difficulties associated with intrinsic disorder, protein instability, or transient interactions, which are inherent in protein structures, are avoided by sequence-based methods, offering a truly revolutionary opportunity to accelerate and create innovative applications in the field of proteomics.

To conclude, even though the *ab initio* reconstruction of PPI networks for model species impressively improved with the years, augmenting the accuracy of the inference within a species remains a problem, especially if we wish to tackle questions on specialized interactions of a protein in different tissues, or differentiating the interaction of paralogous proteins within the organism. For this, one expects to improve in this direction through the development of deep learning architectures that will help us to interpret and disentangle interaction signals better than now. Also, searching for protein interactions in a species without sufficient training information, and training in a model species that is phylogenetically very far away from the species used in testing are still difficult tasks. Testing on multiple species remains a wide-open problem. This is clearly to state that the PPI network reconstruction problem, even when we consider proteins that do not undergo post-transcriptional changes, remains far from being solved but that current achievements can, nonetheless, be applied to the screening of large sets of proteins in search of their partners for many biological questions.

In conclusion, `SENSE-PPI` is a deep learning model that provides results that were not conceivable a few years ago, with an accuracy that could never be reached by state-of-the-art molecular docking approaches applied to proteins with known structures nor deep learning methods before it.

## 4 Materials and Methods

### 4.1 Datasets

#### 4.1.1 Guo’s yeast dataset: a dataset with proteins shared in training and testing

The Guo dataset [12] comprises 2,497 proteins and 11,188 interactions, with an equal number of positive and negative samples. The 5,943 positive interactions were extracted from the DIP 20070219 database [45]. Sequences are more than 50 amino acids long and the majority of them exhibit pairwise sequence identities *<* 40% [12]. The negative samples were generated through a random pairing of proteins that appeared in the positive data but lacked any evidence of interaction. Pairs were subsequently filtered based on the subcellular localization of the proteins, thereby excluding non-interacting pairs residing in the same location. No further filtering was realized in this dataset to allow a fair comparison with models already evaluated on it.

#### 4.1.2 The D-SCRIPT datasets from human and other species

A large human dataset was constructed to test the `D-SCRIPT` deep learning model for the prediction of PPIs [53]. The positive interactions were extracted from STRING (version 11) [56], a database encompassing diverse PPI networks and consolidating comprehensive information from various primary sources. They are high-confidence interactions supported by a positive experimental evidence score in STRING. Protein sequences are greater than 50 and smaller than 800 amino acids in length, and protein pairs exhibit pairwise sequence identities *<* 40%. By construction, any two protein pairs A-B and C-D in the dataset are *non-redundant*, that is both pairs of sequences A, C, and B, D have *<* 40% sequence identity respectively. Eliminating redundancy is crucial as it prevents the model from relying solely on sequence similarity when predicting interactions. To establish an appropriate balance between positive and negative PPIs, negative interactions have been generated by randomly picking two proteins from distinguished pairs in the non-redundant set of positive interactions. These negative samples were created in a 1:10 positive-negative ratio with the aim of better approximating the frequency of positive interactions observed in nature. These construction criteria lead to a primary dataset consisting of 47,932 positive protein interactions and 479,320 negative protein interactions for *H. sapiens*. These interactions were divided into training and testing sets using an 80% - 20% ratio, respectively. Additionally, we have used 4 smaller datasets of *M. musculus, D. melanogaster, C. elegans* and *S. cerevisiae* from [53] made using the procedure described above and containing 5,000 positive and 50,000 negative interactions each.

#### 4.1.3 The SENSE-PPI human dataset

We constructed a second human PPI dataset directly from the STRING database (version 11.5) [58] by focusing exclusively on physical interactions from *H. sapiens*. Similar to the `D-SCRIPT` dataset, we only considered proteins in a length range from 50 to 800 amino acids in order to optimize computational resources and to avoid potential biases introduced by very short or very long sequences. A STRING confidence score threshold of 0.5 was applied and interactions obtained by homology transfer were excluded. We also removed redundant interactions through a clustering strategy that first uses MMseqs2 to cluster proteins at a 40% sequence identity, and then that eliminates redundant pairs as done for the `D-SCRIPT` dataset. The most important difference between the human `D-SCRIPT` dataset and the human `SENSE-PPI` dataset relies on a more stringent condition for constructing negative pairs for training, named here the “neighboring-exclusion” condition. Indeed, a second round of clustering is performed on the proteins in the non-redundant dataset using a 40% sequence identity threshold. Then, a random pair of proteins, A-B, is added to the negative set if no interaction between A and B is known in STRING, and B does not share its cluster with any known interactors of A. This condition ensures that negative pairs are chosen both on the basis of the absence of known interactions and by minimizing the potential biases introduced by homology in shared clusters. As above, in order to reflect the natural frequency of interactions observed in biological systems, the positive-to-negative ratio in the number of protein pairs was set to 1:10. This ratio aims to approximate the relative scarcity of true positive interactions, thus ensuring a more representative distribution of positive and negative examples in the dataset. Overall, the dataset contains 86,304 positive and 863,040 negative interactions.

#### 4.1.4 Ten SENSE-PPI datasets from model and non-model organisms: training on a species and testing on others

We used the same methodology employed for the construction of the `SENSE-PPI` human dataset to build test sets on four non-model organisms, *E. caballus, B. taurus, N. scutatus* and *A. pisum*, and six model organisms, *M. musculus, D. melanogaster, C. elegans, Gallus gallus, S. cerevisiae* and *E. coli*. All these test sets contain 15,000 positive and 150,000 negative entries that were extracted from the STRING v12.0 [57]. In order to conduct additional testing on the non-model organism *A. pisum*, we have also constructed a separate training dataset by merging data from *D. melanogaster* and *C. elegans*. This dataset contains 23,324 positive and 233,240 negative interactions coming from *D. melanogaster* and 17,835 positive and 178,350 negative interactions from *C. elegans*, extracted from STRING v12.0.

#### 4.1.5 The IDPpi dataset describing the interactions between structured proteins and IDPs

The IDPpi dataset [39] contains interactions with intrinsically disordered proteins (IDPs). It is organized in five distinguished datasets corresponding to the testing sets in [39]. Each subset comprises 3,500-4,000 pairs of proteins from *H. sapiens* where one of the partners is intrinsically disordered. Pairs were extracted from the Human Integrated Protein-Protein Interaction rEference (HIPPIE) database [47]. Negative interactions were defined through random sampling: two proteins (one that is an IDP and one that has a stable structure) were selected randomly and were assumed to be non-interacting unless they were already known as interacting. The positive-to-negative ratio of each testing set is 1:1.

#### 4.1.6 Human-Virus PPI dataset

This dataset [61] was designed in order to predict interactions between *H. sapiens* and different viruses. Interacting protein pairs were extracted from the Human-Virus Protein-Protein Interactions database (HVPPI) [65] and negative pairs were constructed using the protocol of the dissimilarity negative sampling, which uses a sequence similarity-based method to explore protein pairs that are unlikely to interact [61].

The training set includes 7,371 positive and 117,326 negative interactions between human and viral proteins and contains 16,933 human and 676 viral proteins in total. Viruses span over many families, and the full list of species included in training can be found in the `SENSE-PPI` repository (`data/human virus` folder).

We constructed two test sets. The first set consists of interactions involving proteins from the *Epstein-Barr* virus. This set encompasses 99 viral proteins and 9,426 human proteins, of which only 596 human proteins demonstrate interaction with *Epstein-Barr* proteins. The set contains a total of 1,308 positive interactions and 14,428 negative interactions. The second dataset exclusively comprises interactions featuring proteins associated with *Influenza* viruses of type A, B, and C. This dataset includes 4,080 positive pairs and 15,724 negative pairs, incorporating a total of 95 viral proteins and 10,594 human proteins. Among these, 1,533 human proteins are involved in interactions with *Influenza* virus proteins. Notably, all proteins from both *Influenza* and *Epstein-Barr* viruses are excluded from the training process. However, it’s important to mention that all human proteins present in the test data have been included in the training set at least once.

**Sequence identity scores for proteins in testing sets**. Given a pair of proteins in a testing set, we define the ‘mean pair sequence identity’ as the mean of the sequence identities of the two proteins relative to the training set. Note that for each protein, we consider the maximum of all sequence identity values between the protein and all proteins in the training set. The computations were performed using the `mmseqs search` command of the MMseqs2 suite [54]. In this particular case, in order to calculate the sequence identity MMseqs2 calculates the ratio of identical aligned residues to the total number of aligned columns, which includes columns containing a gap in either sequence. This can be achieved by using `--alignment-mode 3`. If the search fails to find the similarity with a protein in the training set, the sequence identity value is set to 0 by default.

**Evaluation measures**. To properly evaluate `SENSE-PPI` performance we rely on a ground truth set by reference databases such as the STRING database and the following quantities: known interactions that are identified by `SENSE-PPI` (true positives, TP), interactions identified by `SENSE-PPI` which are not known (false positives, FP), interactions that are not found by `SENSE-PPI` but are known (false negatives, FN), and interactions that are not known and are not detected by `SENSE-PPI` (true negatives, TN). We use four standard measures of performance: recall (sensitivity) = *TP/*(*TP* + *FN*), precision (positive predictive value) = *TP/*(*TP* + *FP*), F1-Score = 2 *TP/*(2 *TP* + *FP* + *FN*), and the Matthews’ correlation coefficient MCC = (*TP · TN − FP · FN*)*/K* where 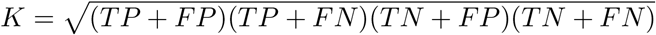.

We also use the precision-recall metric showing the tradeoff between precision and recall for different thresholds. A high area under the curve (AUPRC) represents both high recall and high precision, where high precision relates to a low false positive rate, and high recall relates to a low false negative rate. High scores for both show that the classifier is returning accurate results (high precision), as well as returning a majority of all positive results (high recall). Another measure is the AUROC, calculated as the area under the receiver operating characteristic (ROC) curve, showing the trade-off between true positive rate (TPR) and false positive rate (FPR) across different decision thresholds.

### 4.2 Design of the SENSE-PPI architecture

Recent advancements in deep learning for protein science have lead to the introduction of pre-trained language models [33], which have been developed for natural language processing to learn compressed, informed, and abstract data representations that are later transferred to proteins by adapting them to amino acid sequences. PLMs have demonstrated tremendous potential for a broad range of protein-related problems, such as predicting 3D shapes, interactions, mutational outcomes, and subcellular localizations [42, 1, 15, 40]. They have been trained on large amounts of sequence data, such as the UniProt database [6]. Here, we use ESM2 [25] which is a state-of-the-art general-purpose PLM that has been challenged to predict structure, contact sites, and other protein properties. We used the ESM2 version with 3B parameters, trained on the UniRef50 dataset.

`SENSE-PPI` is designed as a Siamese architecture (Figure 4) consisting of two identical modules with shared weights that are merged together to provide a single output value *p 2* [0, 1] describing the confidence in the interaction, where 0 and 1 stand for low and high confidence respectively. The Siamese design is needed to ensure the commutativity of the model: the output score has to be the same for pairs A-B and B-A.

**Figure 4:**
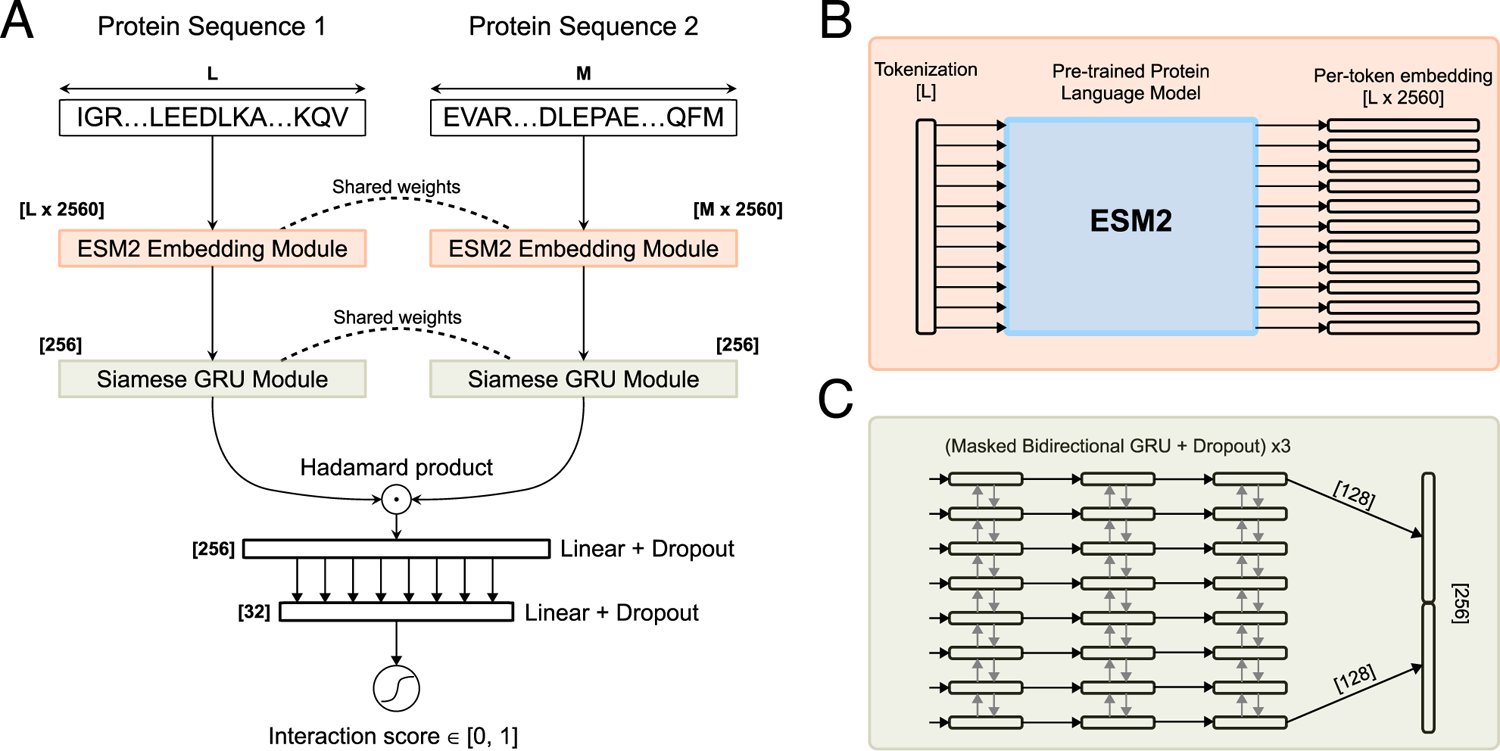
Schematic representation of SENSE-PPI. **A**. `SENSE-PPI` takes two input sequences of lengths L and M through the ESM2 Embedding Module (orange) and transforms them into two large tensors of size *L ×* 2560 and *M ×* 2560 respectively. The two matrices are then independently processed by the Siamese GRU Module (green). The final output vectors are combined together using the Hadamard product and further processed with two linear layers in order to calculate the final interaction score in the interval [0, 1]. **B**. Details of the ESM2 Embedding Module mapping each amino acid in the input sequence into a vector of length 2560. The module is essentially the ESM2 model which comprises 36 layers (3B parameters version) The output of the module is a per-token representation of the input. **C**. Details of the Siamese GRU module composed of three bidirectional GRU layers. The last GRU layer follows a “many-to-one” output scheme that produces a vector of a unified length of 256 (128 for one direction and 128 for the opposite one) for proteins of any length. Since all the sequences are padded to the same fixed size during training every GRU layer was implemented to be dynamical. The layers were configured to accept a sequence mask as a secondary input: this was done in order to process positions in a sequence that belong exclusively to the input data and skip all the padding.

The model takes two amino acid sequences of variable size (*L, M*) as input. Sequences longer than 800 amino acids are trimmed accordingly to this maximum size. The 36 layers of the PLM ESM2 are used to define two embeddings, of size *L* × 2560 and *M* × 2560, for the two amino acid sequences, respectively. Each sequence is processed by one of the Siamese modules sharing the weights with the second module. The Siamese module is composed of 3 layers of gated recurrent units (GRUs) which process the input sequence from one end to another. Each GRU layer is bidirectional, where half of the units are used for one direction of processing, and the other half for the opposite direction. The final layer produces a vector of shape 256 - it takes only the last output of GRU to perform a many-to-one type of processing, where a sequence is “projected” to a single number for each unit. The gated recurrent units process sequences that are masked beforehand: even though all the sequences are padded to the same length to fit the model, the GRU processes only the part that contains the actual sequence and skips the padding. After the GRU layers, the two output vectors are combined together via the Hadamard product. The resulting vector is passed through two linear layers in order to compute the final score. A dropout of 0.5 is applied at every layer (GRU and linear) except for the ESM2 modules.

### 4.3 Other deep learning architectures used for comparison

We conducted a comparative analysis between `SENSE-PPI` and several existing sequence-based deep learning models, namely `PIPR` [4], `D-SCRIPT` [53], `Topsy-Turvy` [52] and `STEP` [28].

`PIPR` is an end-to-end Siamese architecture that incorporates a deep residual recurrent convolutional neural network. This architecture integrates multiple occurrences of convolution layers and residual gated recurrent units. Each amino acid is represented by an embedding that captures its contextual and physico-chemical relationship within the sequence. `PIPR` uses a multigranular feature aggregation process, effectively leveraging the sequential and robust local information present in protein sequences.

`D-SCRIPT` is a deep learning method specifically designed to predict physical PPIs. It features an LSTM-based PLM model for sequence embedding. This is followed by the computation of an outer product between two vector representations, resulting in a three-dimensional tensor. Linear and convolutional layers are then applied to preprocess the tensor, enabling a contact map to be predicted and the probability of interaction to be deduced. `D-SCRIPT` addresses the challenges associated with the weak cross-species generalization exhibited by previous state-of-the-art models. In addition, it establishes a significant overlap with contact maps for protein complexes with known 3D structures.

The `Topsy-Turvy` model combines a traditional sequence-based approach (`D-SCRIPT`) with a network-based approach (GLIDE [7]) for PPI inference. While relying exclusively on sequence data for predictions, `Topsy-Turvy` employs a transfer-learning strategy during training, assimilating insights from global and molecular views of protein interactions. The model achieves state-of-the-art results, enabling genome-scale PPI predictions. It has been applied to non-model organisms.

`STEP` has been developed primarily for the prediction of virus–host PPIs. The core component of the `STEP` architecture is a BERT-based PLM, which generates sequence embeddings. These embeddings are combined through elementwise multiplication and further processed by linear layers.

### 4.4 Implementation details

The implementation of `SENSE-PPI` was carried out using Python 3 and the PyTorch deep learning framework [36]. We trained two versions of the model: a light version, used only for intermediate testing (see Fig. 1E), and a regular version that produced all other results presented here. The regular version is presented in two final forms: the one that was trained on the `D-SCRIPT` human dataset and the one that was trained on the `SENSE-PPI` human dataset. The light version of `SENSE-PPI` has a smaller model size, with an ESM2 embedding block containing 35M parameters and a classification head of 1M parameters. In contrast, the regular version of `SENSE-PPI` has an ESM2 embedding block comprising 3B parameters and a classification head of 2M parameters. The ESM2 module is not retrained, with the only trainable component being the classification head of the model. Each GRU layer contains 256 units followed by a dropout of 0.5. The three final linear layers have 256, 32, and 1 neurons respectively. The training was performed using AdamW optimizer, a batch size of 32, and a learning rate of 10^−4^ with the exception of the Guo’s yeast dataset, for which we used a batch size of 64 and a learning rate of 10^−3^.

The model was trained on a single NVIDIA GeForce RTX 3090 GPU, and on this setup, the training process with the `SENSE-PPI` human dataset, of approximately 1M interactions, for the regular version took approximately five days to complete.

### 4.5 Software availability

`SENSE-PPI` is available at http://gitlab.lcqb.upmc.fr/Konstvv/SENSE-PPI as a package with 5 primary commands: “train”, “test”, “predict”, “predict_string”, and “create_dataset”. The first three commands are used to train the model, obtain metrics on test data, and perform predictions respectively.

The “predict string” command can be used to calculate the predictions for a protein and its known partners according to STRING. The script takes a given number of proteins that are known to interact with a protein of interest, then it performs all versus all predictions and returns the results along with the visualization where one can easily compare the model’s score with real data in STRING. Multiple proteins of interest can be used as input at the same time.

The “create dataset” command creates a dataset for a given species from STRING which is based on the algorithm described in Section “The `SENSE-PPI` human dataset”. The taxon ID is used here as the main input.

The package also contains an internal script to compute ESM2 embeddings. This script is a modification of the original code found in the ESM repository [25], it was slightly edited in order to automate processes during training and testing with `SENSE-PPI`. All datasets used in this study are also provided, including both those newly created and those taken from other publications.

The two trained versions of `SENSE-PPI`, on the `D-SCRIPT` and `SENSE-PPI` human datasets, are available in the `SENSE-PPI` repository in the `pretrained_models` folder, under the names `dscript.ckpt` and `senseppi.ckpt` respectively. In addition, this folder also contains pretrained versions used for other tests presented in this work such as the human–virus model, and models based on a combination of model species (fly/worm as well as fly/worm/human).

## Acknowledgments

Bio initiative at Sorbonne University for the PhD fellowship and travel grants to KV; Sorbonne Center for Artificial Intelligence (SCAI) and LIP6 laboratory (UMR 7606, Sorbonne University-CNRS) for access to their clusters; Agence Nationale de Recherches sur le Sida et les Hepatites Virales [ANRS–AAP-2021-CSS-12] (AC).

## Notes

http://gitlab.lcqb.upmc.fr/Konstvv/SENSE-PPI

### Competing Interest Statement

The authors have declared no competing interest.

### Summary of Updates

Figure 4 revised (fixed typo in embedding dimensionality)

## References

[1] E. C. Alley, G. Khimulya, S. Biswas, M. AlQuraishi, and G. M. Church. Unified rational protein engineering with sequence-based deep representation learning. Nature methods, 16(12):1315–1322, 2019.

[2] A. Bensimon, A. J. Heck, and R. Aebersold. Mass spectrometry-based proteomics and network biology. Annu. Rev. Biochem., 81:379, 2012.

[3] D. Carroll, P. Daszak, N. D. Wolfe, G. F. Gao, C. M. Morel, S. Morzaria, A. Pablos-Méndez, O. Tomori, and J. A. Mazet. The global virome project. Science, 359(6378):872–874, 2018.

[4] M. Chen, C. J. T. Ju, G. Zhou, X. Chen, T. Zhang, K.-W. Chang, C. Zaniolo, and W. Wang. Multifaceted protein–protein interaction prediction based on Siamese residual RCNN. Bioinformatics, 35(14):i305–i314, 07 2019.

[5] K. Cho, B. van Merriënboer, D. Bahdanau, and Y. Bengio. On the properties of neural machine translation: Encoder–decoder approaches. In Proceedings of SSST-8, Eighth Workshop on Syntax, Semantics and Structure in Statistical Translation, pages 103–111, Doha, Qatar, Oct. 2014. Association for Computational Linguistics.

[6] T. U. Consortium. UniProt: the universal protein knowledgebase in 2021. Nucleic Acids Research, 49(D1):D480–D489, 11 2020.

[7] K. Devkota, J. M. Murphy, and L. J. Cowen. GLIDE: combining local methods and diffusion state embeddings to predict missing interactions in biological networks. Bioinformatics, 36(Supplement 1):i464–i473, 07 2020.

[8] A. Elnaggar, M. Heinzinger, C. Dallago, G. Rehawi, Y. Wang, L. Jones, T. Gibbs, T. Feher, C. Angerer, M. Steinegger, D. Bhowmik, and B. Rost. Prottrans: Towards cracking the language of life’s code through self-supervised deep learning and high performance computing. bioRxiv, 2020.

[9] E. L. Folador, S. S. Hassan, N. Lemke, D. Barh, A. Silva, R. S. Ferreira, and V. Azevedo. An improved interolog mapping-based computational prediction of protein-protein interactions with increased network coverage. Integr. Biol. (Camb.), 6:1080, 2014.

[10] J. Garcia-Garcia, S. Schleker, J. Klein-Seetharaman, and B. Oliva. Bips: Biana interolog prediction server. a tool for protein-protein interaction inference. Nucleic Acids Res., 40:W147, 2012.

[11] T. Guirimand, S. Delmotte, and V. Navratil. Virhostnet 2.0: surfing on the web of virus/host molecular interactions data. Nucleic acids research, 43(D1):D583–D587, 2015.

[12] Y. Guo, L. Yu, Z. Wen, and M. Li. Using support vector machine combined with auto covariance to predict protein–protein interactions from protein sequences. Nucleic Acids Research, 36(9):3025–3030, 04 2008.

[13] T. Hamp and B. Rost. Evolutionary profiles improve protein–protein interaction prediction from sequence. Bioinformatics, 31(12):1945–1950, 02 2015.

[14] G. T. Hart, A. K. Ramani, and E. M. Marcotte. How complete are current yeast and human protein-interaction networks? Genome Biology, 7(11):120, 2006.

[15] M. Heinzinger, A. Elnaggar, Y. Wang, C. Dallago, D. Nechaev, F. Matthes, and B. Rost. Modeling aspects of the language of life through transfer-learning protein sequences. BMC bioinformatics, 20(1):1–17, 2019.

[16] M. Huynen, B. Snel, W. Lathe, 3rd, and P. Bork. Predicting protein function by genomic context: quantitative evaluation and qualitative inferences. Genome Res, 10(8):1204–10, Aug 2000.

[17] T. Ito, T. Chiba, R. Ozawa, M. Yoshida, M. Hattori, and Y. Sakaki. A comprehensive two-hybrid analysis to explore the yeast protein interactome. Proc. Natl. Acad. Sci. U. S. A., 98:4569, 2001.

[18] S. Kumar, G. Stecher, M. Suleski, and S. B. Hedges. Timetree: a resource for timelines, timetrees, and divergence times. Molecular biology and evolution, 34(7):1812–1819, 2017.

[19] A. Laddach, S. S. Chung, and F. Fraternali. Prediction of protein-protein interactions: Looking through the kaleidoscope. In Encyclopedia of Bioinformatics and Computational Biology: ABC of Bioinformatics, pages 834–848. Elsevier, 2018.

[20] A. Laddach, J. C.-F. Ng, S. S. Chung, and F. Fraternali. Genetic variants and protein–protein interactions: a multidimensional network-centric view. Current opinion in structural biology, 50:82–90, 2018.

[21] E. Laine and A. Carbone. Protein social behavior makes a stronger signal for partner identification than surface geometry. Proteins: Structure, Function, and Bioinformatics, 85(1):137–154, 2017.

[22] E. D. Levy, C. R. Landry, and S. W. Michnick. How perfect can protein interactomes be? Science signaling, 2(60):pe11–pe11, 2009.

[23] H. A. Lewin, G. E. Robinson, W. J. Kress, W. J. Baker, J. Coddington, K. A. Crandall, R. Durbin, S. V. Edwards, F. Forest, M. T. P. Gilbert, et al. Earth biogenome project: Sequencing life for the future of life. Proceedings of the National Academy of Sciences, 115(17):4325–4333, 2018.

[24] F. Li, X. Zhao, M. Li, K. He, C. Huang, Y. Zhou, Z. Li, and J. R. Walters. Insect genomes: progress and challenges. Insect molecular biology, 28(6):739–758, 2019.

[25] Z. Lin, H. Akin, R. Rao, B. Hie, Z. Zhu, W. Lu, N. Smetanin, R. Verkuil, O. Kabeli, Y. Shmueli, A. dos Santos Costa, M. Fazel-Zarandi, T. Sercu, S. Candido, and A. Rives. Evolutionary-scale prediction of atomic level protein structure with a language model. bioRxiv, 2022.

[26] B. T. Lobingier, R. Hüttenhain, K. Eichel, K. B. Miller, A. Y. Ting, M. von Zastrow, and N. J. Krogan. An approach to spatiotemporally resolve protein interaction networks in living cells. Cell, 169(2):350–360, 2017.

[27] A. Lopes, S. Sacquin-Mora, V. Dimitrova, E. Laine, Y. Ponty, and A. Carbone. Protein-protein interactions in a crowded environment: an analysis via cross-docking simulations and evolutionary information. PLoS Comput Biol, 9(12):e1003369, 2013.

[28] S. Madan, V. Demina, M. Stapf, O. Ernst, and H. Frohlich. Accurate prediction of virus-host protein-protein interactions via a siamese neural network using deep protein sequence embeddings. Patterns, 3(9):100551, 2022.

[29] L. R. Matthews, P. Vaglio, J. Reboul, H. Ge, B. P. Davis, J. Garrels, S. Vincent, and M. Vidal. Identification of potential interaction networks using sequence-based searches for conserved protein-protein interactions or “interologs”. Genome Res, 11(12):2120–6, Dec 2001.

[30] I. Morilla, J. G. Lees, A. J. Reid, C. Orengo, and J. A. Ranea. Assessment of protein domain fusions in human protein interaction networks prediction: Application to the human kinetochore model. New Biotechnol., 27:755, 2010.

[31] R. Mosca, A. Céol, and P. Aloy. Interactome3d: adding structural details to protein networks. Nature methods, 10(1):47, 2013.

[32] M. E. Oates, P. Romero, T. Ishida, M. Ghalwash, M. J. Mizianty, B. Xue, Z. Dosztányi, V. N. Uversky, Z. Obradovic, L. Kurgan, et al. D2p2: database of disordered protein predictions. Nucleic acids research, 41(D1):D508–D516, 2012.

[33] D. Ofer, N. Brandes, and M. Linial. The language of proteins: Nlp, machine learning & protein sequences. Computational and Structural Biotechnology Journal, 19:1750–1758, 2021.

[34] S. Orchard, M. Ammari, B. Aranda, L. Breuza, L. Briganti, F. Broackes-Carter, N. H. Campbell, G. Chavali, C. Chen, N. Del-Toro, et al. The mintact project—intact as a common curation platform for 11 molecular interaction databases. Nucleic acids research, 42(D1):D358–D363, 2014.

[35] Y. Park and E. M. Marcotte. Flaws in evaluation schemes for pair-input computational predictions. Nature methods, 9(12):1134, 2012.

[36] A. Paszke, S. Gross, F. Massa, A. Lerer, J. Bradbury, G. Chanan, T. Killeen, Z. Lin, N. Gimelshein, L. Antiga, A. Desmaison, A. Kopf, E. Yang, Z. DeVito, M. Raison, A. Tejani, S. Chilamkurthy, B. Steiner, L. Fang, J. Bai, and S. Chintala. Pytorch: An imperative style, high-performance deep learning library. In Advances in Neural Information Processing Systems 32, pages 8024–8035. Curran Associates, Inc., 2019.

[37] M. Pellegrini, E. M. Marcotte, M. J. Thompson, D. Eisenberg, and T. O. Yeates. Assigning protein functions by comparative genome analysis: Protein phylogenetic profiles. Proc. Natl. Acad. Sci. U. S. A., 96:4285, 1999.

[38] M. M. Pentony and D. T. Jones. Modularity of intrinsic disorder in the human proteome. Proteins: Structure, Function, and Bioinformatics, 78(1):212–221, 2010.

[39] V. Perovic, N. Sumonja, L. A. Marsh, S. Radovanovic, M. Vukicevic, S. G. E. Roberts, and N. Veljkovic. Idppi: Protein-protein interaction analyses of human intrinsically disordered proteins. Scientific Reports, 10563(8), 2018.

[40] R. Rao, N. Bhattacharya, N. Thomas, Y. Duan, P. Chen, J. Canny, P. Abbeel, and Y. Song. Evaluating protein transfer learning with tape. Advances in neural information processing systems, 32, 2019.

[41] A. Rhie, S. A. McCarthy, O. Fedrigo, J. Damas, G. Formenti, S. Koren, M. Uliano-Silva, W. Chow, A. Fungtammasan, J. Kim, et al. Towards complete and error-free genome assemblies of all vertebrate species. Nature, 592(7856):737–746, 2021.

[42] A. Rives, J. Meier, T. Sercu, S. Goyal, Z. Lin, J. Liu, D. Guo, M. Ott, C. L. Zitnick, J. Ma, et al. Biological structure and function emerge from scaling unsupervised learning to 250 million protein sequences. Proceedings of the National Academy of Sciences, 118(15):e2016239118, 2021.

[43] T. Rolland, M. Taşan, B. Charloteaux, S. J. Pevzner, Q. Zhong, N. Sahni, S. Yi, I. Lemmens, C. Fontanillo, R. Mosca, et al. A proteome-scale map of the human interactome network. Cell, 159(5):1212–1226, 2014.

[44] S. Sacquin-Mora, A. Carbone, and R. Lavery. Identification of protein interaction partners and protein– protein interaction sites. Journal of molecular biology, 382(5):1276–1289, 2008.

[45] L. Salwinski, C. S. Miller, A. J. Smith, F. K. Pettit, J. U. Bowie, and D. Eisenberg. The database of interacting proteins: 2004 update. Nucleic acids research, 32(Suppl. 1):D449–D451, 2004.

[46] M. H. Schaefer, J.-F. Fontaine, A. Vinayagam, P. Porras, E. E. Wanker, and M. A. Andrade-Navarro. Hippie: Integrating protein interaction networks with experiment based quality scores. PloS one, 7(2):e31826, 2012.

[47] M. H. Schaefer, J. F. Fontaine, A. Vinayagam, P. Porras, E. E. Wanker, and M. A. Andrade-Navarro. Hippie: Integrating protein interaction networks with experiment based quality scores. PLoS One, 7:e31826, 2012.

[48] J. D. Scott and T. Pawson. Cell signaling in space and time: where proteins come together and when they’re apart. Science, 326(5957):1220–1224, 2009.

[49] M. S. Scott and G. J. Barton. Probabilistic prediction and ranking of human protein-protein interactions. BMC Bioinformatics, 8:239, Jul 2007.

[50] B. Seoane and A. Carbone. The complexity of protein interactions unravelled from structural disorder. PLOS Computational Biology, 17(1):e1008546, 2021.

[51] B. Seoane and A. Carbone. Soft disorder modulates the assembly path of protein complexes. PLOS Computational Biology, 18(11):e1010713, 2022.

[52] R. Singh, K. Devkota, S. Sledzieski, B. Berger, and L. Cowen. Topsy-Turvy: integrating a global view into sequence-based PPI prediction. Bioinformatics, 38(Supplement 1):i264–i272, 06 2022.

[53] S. Sledzieski, R. Singh, L. Cowen, and B. Berger. D-script translates genome to phenome with sequence-based, structure-aware, genome-scale predictions of protein-protein interactions. Cell Systems, 12(10), 2021.

[54] M. Steinegger and J. Söding. Mmseqs2 enables sensitive protein sequence searching for the analysis of massive data sets. Nature Biotechnology, 35(11):1026–1028, Oct 2017.

[55] M. P. H. Stumpf, T. Thorne, E. de Silva, R. Stewart, H. J. An, M. Lappe, and C. Wiuf. Estimating the size of the human interactome. Proc Natl Acad Sci U S A, 105(19):6959–64, May 2008.

[56] D. Szklarczyk, A. L. Gable, D. Lyon, A. Junge, S. Wyder, J. Huerta-Cepas, M. Simonovic, N. T. Doncheva, J. H. Morris, P. Bork, L. J. Jensen, and C. Mering. STRING v11: protein–protein association networks with increased coverage, supporting functional discovery in genome-wide experimental datasets. Nucleic Acids Research, 47(D1):D607–D613, 11 2018.

[57] D. Szklarczyk, A. L. Gable, K. C. Nastou, D. Lyon, R. Kirsch, S. Pyysalo, N. T. Doncheva, M. Legeay, T. Fang, P. Bork, L. J. Jensen, and C. von Mering. The STRING database in 2021: customizable protein–protein networks, and functional characterization of user-uploaded gene/measurement sets. Nucleic Acids Research, 49(D1):D605–D612, 11 2020.

[58] D. Szklarczyk, R. Kirsch, M. Koutrouli, K. Nastou, F. Mehryary, R. Hachilif, A. L. Gable, T. Fang, N. Doncheva, S. Pyysalo, P. Bork, L. Jensen, and C. von Mering. The STRING database in 2023: protein–protein association networks and functional enrichment analyses for any sequenced genome of interest. Nucleic Acids Research, 51(D1):D638–D646, 11 2022.

[59] D. Szklarczyk, J. H. Morris, H. Cook, M. Kuhn, S. Wyder, M. Simonovic, A. Santos, N. T. Doncheva, A. Roth, P. Bork, et al. The string database in 2017: quality-controlled protein–protein association networks, made broadly accessible. Nucleic acids research, page gkw937, 2016.

[60] L. R. Thompson, J. G. Sanders, D. McDonald, A. Amir, J. Ladau, K. J. Locey, R. J. Prill, A. Tripathi, S. M. Gibbons, G. Ackermann, et al. A communal catalogue reveals earth’s multiscale microbial diversity. Nature, 551(7681):457–463, 2017.

[61] S. Tsukiyama, M. M. Hasan, S. Fujii, and H. Kurata. Lstm-phv: prediction of human-virus protein–protein interactions by lstm with word2vec. Briefings in bioinformatics, 22(6):bbab228, 2021.

[62] P. Uetz, L. Giot, G. Cagney, T. A. Mansfield, R. S. Judson, J. R. Knight, D. Lockshon, V. Narayan, M. Srinivasan, P. Pochart, A. Qureshi-Emili, Y. Li, B. Godwin, D. Conover, T. Kalbfleisch, G. Vijayadamodar, M. Yang, M. Johnston, S. Fields, and J. M. Rothberg. A comprehensive analysis of protein-protein interactions in saccharomyces cerevisiae. Nature, 403:623, 2000.

[63] R. Van Der Lee, M. Buljan, B. Lang, R. J. Weatheritt, G. W. Daughdrill, A. K. Dunker, M. Fuxreiter, J. Gough, J. Gsponer, D. T. Jones, et al. Classification of intrinsically disordered regions and proteins. Chemical reviews, 114(13):6589–6631, 2014.

[64] M. N. Wass, G. Fuentes, C. Pons, F. Pazos, and A. Valencia. Towards the prediction of protein interaction partners using physical docking. Molecular systems biology, 7(1):469, 2011.

[65] X. Yang, X. Lian, C. Fu, S. Wuchty, S. Yang, and Z. Zhang. Hvidb: a comprehensive database for human–virus protein–protein interactions. Briefings in bioinformatics, 22(2):832–844, 2021.

[66] H. Yu, P. Braun, M. A. Yildirim, I. Lemmens, K. Venkatesan, J. Sahalie, T. Hirozane-Kishikawa, F. Gebreab, N. Li, and N. Simonis. High-quality binary protein interaction map of the yeast interactome network. Science, 322:104, 2008.

[67] H. Yu, N. M. Luscombe, H. X. Lu, X. Zhu, Y. Xia, J.-D. J. Han, N. Bertin, S. Chung, M. Vidal, and M. Gerstein. Annotation transfer between genomes: protein-protein interologs and protein-dna regulogs. Genome Res, 14(6):1107–18, Jun 2004.

