## Supplementary material for "SENSE-PPI reconstructs protein-protein interactions of various complexities, within, across, and between species, with sequence-based evolutionary scale modeling and deep learning": Tables SI 1-6

### Supplementary Information

| Model | F1-Score | MCC |
|---|---|---|
| Guo et al. [12] | 87.34 ± 1.33 | 75.09 ± 2.51 |
| PIPR [4] | 97.09 ± 0.27 | 94.17 ± 0.48 |
| STEP [28] | 97.37 ± 0.27 | 94.77 ± 0.54 |
| SENSE-PPI | **97.73 ± 0.31** | **95.46 ± 0.61** |

**Table SI 1: Comparative analysis of the performance of SENSE-PPI and several other DL architectures on Guo's yeast dataset.** Comparison is made based on a 5-fold cross-validation test between SENSE-PPI, PIPR, STEP, and the original model from Guo et al. Values reported for all DL architectures other than SENSE-PPI were taken from [28]. The best values across architectures are shown in bold.

| Model | AUROC | AUPRC | MCC | F1-Score |
|---|---|---|---|---|
| SENSE-PPI | **0.751±0.004** | **0.747±0.008** | 0.346±0.011 | 0.578±0.011 |
| IDPpi | 0.746±0.017 | 0.734±0.020 | **0.348±0.028** | – |

**Table SI 2: Performance of SENSE-PPI on interactions with intrinsically disordered proteins.** Performance comparison between SENSE-PPI and IDPpi on the interaction dataset of human intrinsically disordered proteins extracted from [39].

| Test set | AUPRC | AUROC | MCC | F1-Score |
|---|---|---|---|---|
| *Epstein-Barr virus* | 0.799 ± 0.022 | 0.932 ± 0.008 | 0.729 ± 0.023 | 0.738 ± 0.019 |
| *Influenza viruses* | 0.883 ± 0.006 | 0.931 ± 0.006 | 0.736 ± 0.006 | 0.755 ± 0.007 |

**Table SI 3: Evaluation of SENSE-PPI on human-virus interactions.** The test was done on PPIs that belong to Epstein-Barr and Influenza viruses. These viruses were excluded from training, whereas human proteins present in testing were also presented in training.

| Test set | Mean pair sequence identity | AUPRC | AUROC | MCC | F1-Score |
|---|---|---|---|---|---|
| *M. musculus* | 0.829 | 0.877 | 0.970 | 0.784 | 0.804 |
| *B. taurus* | 0.837 | 0.812 | 0.950 | 0.740 | 0.764 |
| *E. caballus* | 0.851 | 0.792 | 0.938 | 0.723 | 0.747 |
| *G. gallus* | 0.715 | 0.823 | 0.941 | 0.738 | 0.762 |
| *N. scutatus* | 0.671 | 0.705 | 0.890 | 0.639 | 0.671 |
| *D. melanogaster* | 0.472 | 0.792 | 0.942 | 0.686 | 0.715 |
| *A. pisum* | 0.447 | 0.704 | 0.886 | 0.626 | 0.660 |
| *C. elegans* | 0.410 | 0.766 | 0.929 | 0.664 | 0.695 |
| *S. cerevisiae* | 0.307 | 0.666 | 0.918 | 0.561 | 0.594 |
| *E. coli* | 0.124 | 0.295 | 0.663 | 0.199 | 0.277 |

**Table SI 4: Evaluation of SENSE-PPI trained on the SENSE-PPI human dataset on model and non-model organisms.** The test was performed on PPIs that belong to 10 different species, 6 from model organisms and 4 from non-model organisms.

| | | AUPRC | AUROC | MCC | F1 Score |
|---|---|---|---|---|---|
| *Influenza viruses* | 1 | 0.876 | 0.925 | 0.733 | 0.754 |
| | 2 | 0.880 | 0.925 | 0.737 | 0.753 |
| | 3 | 0.881 | 0.928 | 0.740 | 0.758 |
| | 4 | 0.890 | 0.936 | 0.726 | 0.742 |
| | 5 | 0.892 | 0.940 | 0.744 | 0.764 |
| | 6 | 0.880 | 0.932 | 0.737 | 0.760 |
| *Epstein-Barr virus* | 1 | 0.815 | 0.938 | 0.747 | 0.757 |
| | 2 | 0.804 | 0.925 | 0.735 | 0.737 |
| | 3 | 0.792 | 0.922 | 0.718 | 0.721 |
| | 4 | 0.825 | 0.945 | 0.751 | 0.757 |
| | 5 | 0.799 | 0.935 | 0.741 | 0.749 |
| | 6 | 0.757 | 0.927 | 0.684 | 0.707 |

**Table SI 5: Evaluation of `SENSE-PPI` on human-virus interactions.** Both statistics for Epstein-Barr and Influenza viruses were computed out of 6 independent training runs. The mean and standard deviation are reported in Table SI 3.

| Testing set | AUPRC | AUROC | MCC | F1 Score |
|---|---|---|---|---|
| 1 | 0.756 | 0.755 | 0.335 | 0.560 |
| 2 | 0.747 | 0.751 | 0.345 | 0.572 |
| 3 | 0.735 | 0.743 | 0.349 | 0.579 |
| 4 | 0.756 | 0.755 | 0.364 | 0.591 |
| 5 | 0.742 | 0.749 | 0.335 | 0.588 |

**Table SI 6: Evaluation of `SENSE-PPI` on the IDPpi dataset.** The test was performed on 5 preprocessed datasets extracted from [39]. The mean and standard deviation for each metric are presented in Table SI 2.
